# Loss of calcium-binding protein *Cbp53E* leads to delayed repolarization of photoreceptor cells in Drosophila

**DOI:** 10.64898/2026.02.13.705836

**Authors:** Natalie Cmejla, Celia Brekken, Justin Chilson, Regan Alexander, Natalie Cleary, Kathryn Davis, Shane Gonnelly, Eric Hawks, Glenn Jordan, Austin Link, Cynthia Ndahr, Gavin Olson, Estevan Quintana, Brandon Schultz, Katherine Scott, Sebastian Spencer, Margaret Talafuse, Jordn Wolsky, Hailey Zwirner, Rodrigo da Costa Aparecido, David S. Ronderos

**Affiliations:** School of Arts and Sciences, University of Mary, 7500 University Dr., Bismarck, ND 58504; Hamm School of Engineering, University of Mary, 7500 University Dr., Bismarck, ND 58504

## Abstract

Calcium functions as an important second messenger in a wide variety of intracellular processes. In photoreceptor cells, calcium is involved in activation, deactivation, and adaptation in response to light stimuli. Calcium-binding protein 53E (Cbp53E, also known as calbindin-32 or cbn), a protein with 6 EF-Hand domains thought to act as a calcium buffer, was previously identified to have elevated expression levels in the eye of drosophila. While a recent study showed that transgenic flies lacking Cbp53E have aberrant axonal arborization at the neuromuscular junction, nothing is known about the role of Cbp53E in the visual system. We performed electroretinogram (ERG) recordings on Cbp53E mutant flies to test whether eye function was affected. Here, we report that Cbp53E null mutants exhibit a prolonged repolarization (or slow termination) phenotype which can be rescued by expressing Cbp53E in photoreceptor cells. The human homologs Calbindin 2, Calbindin 1, and S100G also rescue the Drosophila ERG phenotype. This supports a role for Cbp53E in regulating intracellular calcium levels of photoreceptor cells and contributing to normal sensory neuron response dynamics *in vivo* in Drosophila and suggests a similar function in human photoreceptor cells as well.

## INTRODUCTION

The Drosophila compound eye is comprised of ∼800 units known as ommatidia (reviewed in (Montell, 2012; Wang & Montell, 2007a)). Each ommatidium is made up of multiple cell types, including pigment cells and light-sensitive photoreceptor cells (Montell, 2012; Wang & Montell, 2007a). Light-dependent activation of photoreceptor cells is initiated by the G-protein coupled receptor, rhodopsin (Nichols & Pak, 1985). The activation of rhodopsin triggers a G-protein signaling cascade that ultimately results in the opening of ion channels Trp and Trpl, causing the influx of Ca^2+^ and Na^+^ ions (Montell & Rubin, 1989; Niemeyer et al., 1996). Changes in intracellular Ca^2+^ are known to underlie numerous processes in the photoreceptor cell, including depolarization, repolarization, and adaptation (reviewed in (Voolstra & Huber, 2020)).

Cbp53E was identified to have eye-enriched expression in Drosophila (Xu et al., 2004). The *Cbp53E* gene encodes a protein with six EF-hand domains, and is predicted to function as a calcium buffer (Öztürk-Çolak et al., 2024). Drosophila Cbp53E null mutant flies have been generated and described previously, and were shown to display aberrant developmental patterns in the neuromuscular junction (Hagel et al., 2015). However, the effects of Cbp53E mutation on vision and the role of Cbp53E in the Drosophila visual system is unknown. We therefore set out to determine whether Cbp53E was involved in vision in Drosophila.

## MATERIALS AND METHODS

### Fly Stocks

Control stocks used were *w*^*1118*^ from the Bloomington Drosophila Stock Center (Cook et al., 2010). *Cbp53E*^*mi22*^, *Cbp53E*^*mi41*^, and *UAS-Cbp53E* stocks were a gift from Dr. Charles Tessier (Hagel et al., 2015). The *rdhb-GAL4* and *ninaE-GAL4* stocks were a gift from Dr. Craig Montell (Wang et al., 2012). Genotypes used in rescue experiments were *rdhb-GAL4/+;Cbp53E*^*mi22*^*;UAS-Cbp53E/+* (for pigment cell-specific expression), and *+;Cbp53E*^*mi22*^*;ninaE-GAL4/UAS-Cbp53E* (for photoreceptor cell-specific expression). Plasmids for expression of *UAS-hCalb2* (Vector ID: VB230210-1395bmk), *UAS-hCalb1* (Vector ID: VB230210-1394ndm), and *UAS-hS100G* (Vector ID: VB230210-1396puk) were synthesized by VectorBuilder (Watertown, MA) and injected into *Cbp53E*^*mi22*^ mutant flies by BestGene Inc. (Chino, CA) to generate transgenic flies for rescue experiments.

### Fly Husbandry & Rearing Conditions

Flies were fed on instant fly medium (Genesee) and housed at room temperature in custom built light cycling chambers that used commercial LED light strips (Patriot Lighting) of either white or blue light. Lights were controlled by an outlet timer to deliver 18 hours of white light and 6 hours of darkness per day in Figure 1, Figure 2, and Figure 4. Timers were eliminated to deliver constant blue light in Figure 3.

**Figure 1.**
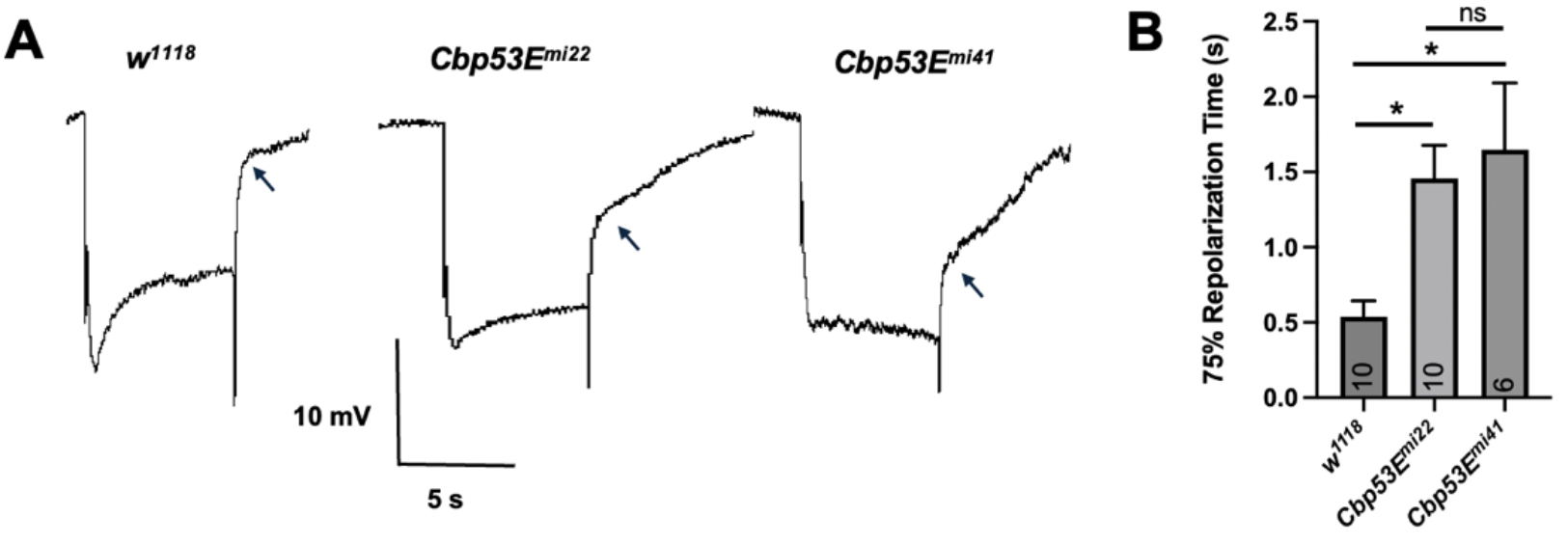
*Cbp53E* null mutant fly ERGs exhibit slow repolarization. **(A)** Sample ERG traces from two independent lines of *Cbp53E* null mutant flies compared to wild-type controls. Arrows indicate observed differences in repolarization characteristics. **(B)** Quantification of ERG traces measuring the time needed to reach 75% of the initial voltage following cessation of light stimulus in flies of the indicated genotype age 1-14 days. Columns represent mean and error bars represent SEM; *n* number is shown within columns, and asterisks (*) indicate results of statistical analysis by One-Way ANOVA followed by Tukey’s pairwise comparison.

**Figure 2.**
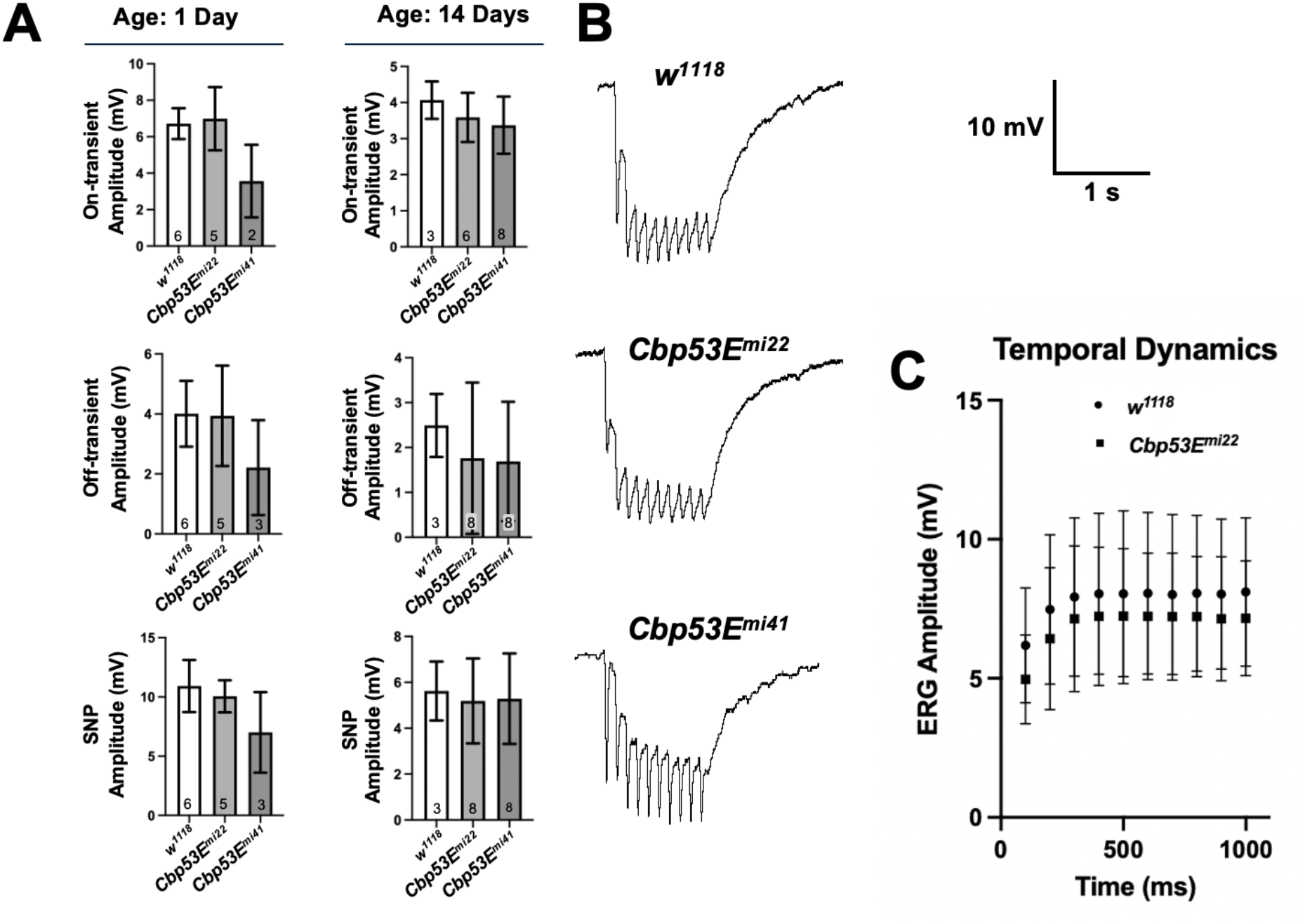
Comparison of other ERG components in *w*^*1118*^ and *Cbp53E* null mutants. Quantification of ERG components **(A)** On-transient, Off-transient, and Sustained Negative Potential (SNP) amplitudes. Columns and error bars represent mean and SEM. *n* numbers are within the columns. **(B)** ERG traces obtained using tetanic stimulation protocol. **(C)** Quantification of of ERG amplitudes shown in panel **B**, represents mean and SEM, *n* = 8 each.

**Figure 3.**
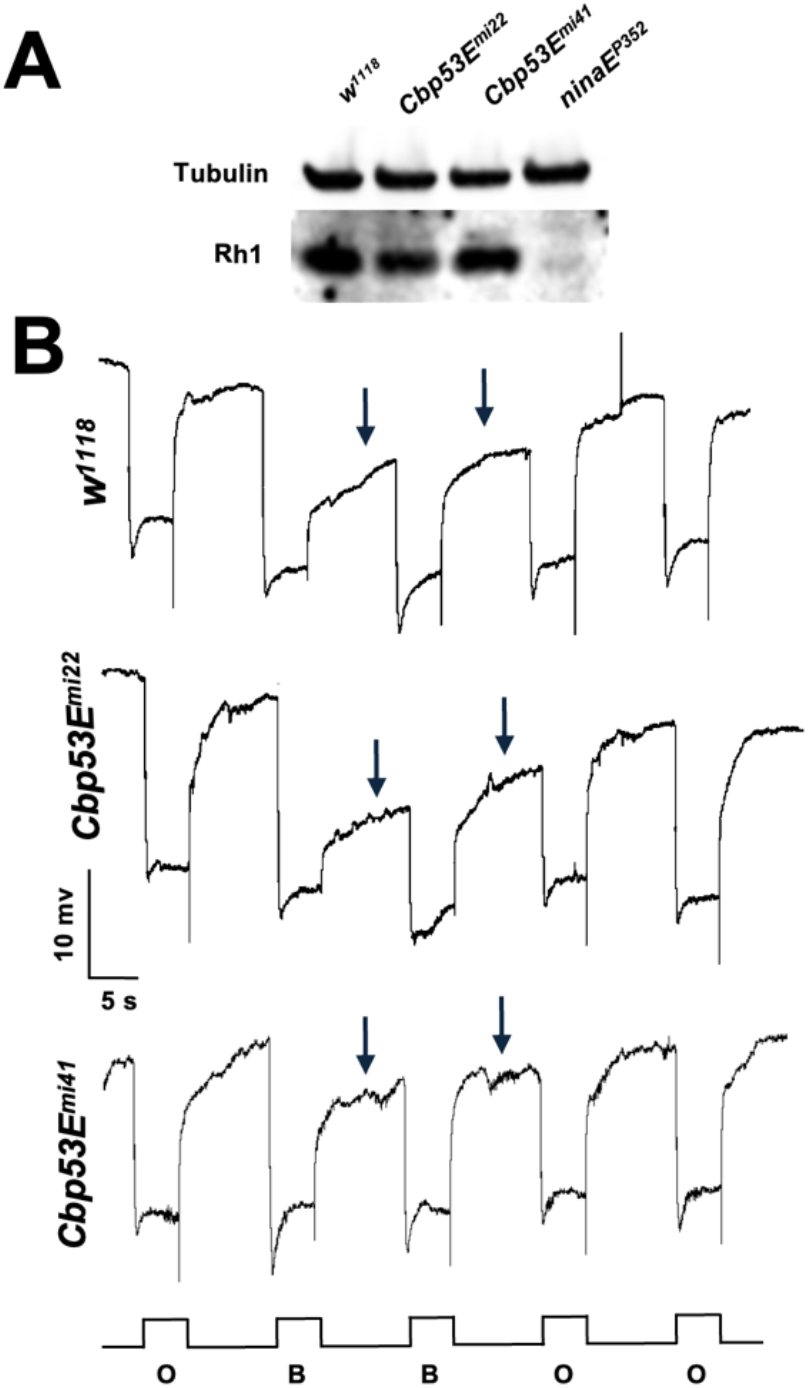
*Cbp53E* mutants do not undergo retinal degeneration. **(B)** Western blot of Rhodopsinl protein for the indicated genotypes. **(C)** ERG traces for the indicated genotypes using a prolonged depolarizing afterpotential (PDA) protocol after 5 days constant blue-light exposure. Light stimulus using either orange (O) or blue (B) light is indicated below each trace. Arrows indicate the PDA.

**Figure 4.**
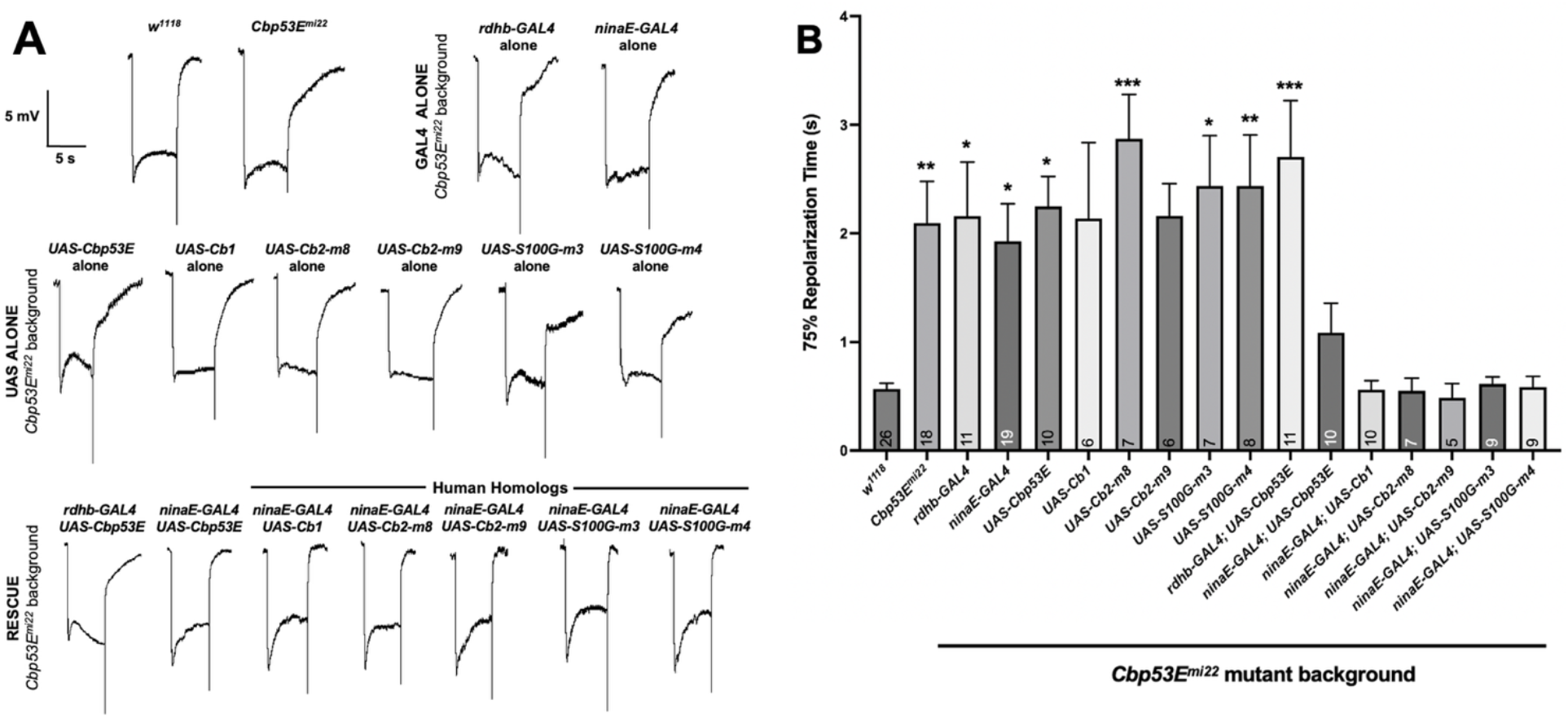
Rescue of *Cbp53E*^*mi22*^ Mutant ERG Phenotype By Misexpression of Drosophila and Human Calcium-binding Proteins in Photoreceptor Cells. **(A)** Sample ERG traces of the indicated genotypes. All GAL4, UAS, and Rescue lines are in a *Cbp53E*^*mi22*^ null mutant background. **(B)** Quantification of ERG results showing mean and SEM, with *n* numbers shown in each column. Asterisks (*) indicate results One-Way ANOVA with post-hoc Tukey’s pairwise comparison with *w*^*1118*^ results.

### Electrophysiology

ERG recordings were performed by placing a recording electrode onto a drop of Signa electrode conducting cream (Parker Laboratories, Inc.) that was applied onto the eye and a reference electrode was placed onto the thorax (for a more detailed explanation of Drosophila ERG recordings, see (Vilinsky & Johnson, 2012)). Recording and reference electrodes were made using a Model P-97 needle puller (Sutter Instruments) and filled with Ringer’s solution. Signals were amplified using an Intracellular Electrometer IE-210 (Warner Instruments) and recorded using a PowerLab 4/30 data acquisition box and LabChart software (ADInstruments). Light stimulus was produced using either a custom-built LED controller or an X-Cite Turbo multi-wavelength LED Illumination system (Excelitas). For the PDA experiments, we used a blue 480nm band pass filter or an orange 580nm band pass filter (Newport). All other light stimuli were white light. Each *n* represents a separate fly, with each fly being tested only once.

### Data Analysis

Data was analyzed in LabChart Reader (ADInstruments). Repolarization times were calculated by taking the initial voltage prior to light activation minus the SNP voltage prior to light deactivation, then multiplying that number by 75% to obtain the 75% repolarization voltage. Then we measure the time from the 75% repolarization voltage to the SNP voltage prior to light deactivation to get the Time to 75% Repolarization shown in the graphs. Flies of similar age, rearing conditions, and genotypes were grouped together and compared to other genotypes as indicated. Statistical analysis was performed in GraphPad Prism using one-way ANOVA and Tukey comparison of the values (GraphPad Prism). Data excluded were either poor quality recordings (e.g., if the signal to noise ratio of the recording was deemed excessive per qualitative judgement) or statistical outliers identified using a ROUT test with a Q value of 1% in (GraphPad Prism).

### Western blot

Western blots were performed using dissected drosophila heads homogenized on ice in Laemmli buffer (Bio-Rad) with added β-mercaptoethanol (Sigma-Aldrich), incubated for 1min. at 40°C in a heat block, and loaded into a 4-20% pre-cast gradient gel (Bio-Rad) and run for 30min. at 100V. Proteins were transferred overnight at 30V to PVDF membrane (Millipore Sigma) and blocked in TBS-T for 1 hour (LI-COR). Primary antibodies diluted in TBS-T against drosophila rhodopsin 1 (4C5) or alpha-tubulin (12G10 anti-alpha-tubulin) were incubated overnight at room temperature (Developmental Studies Hybridoma Bank). Secondary antibodies (Goat anti-mouse IR700 dye, LI-COR) in TBS-T were incubated for 1hr. and then blots visualized and quantified using an LI-COR Odyssey Fc Imager (LI-COR).

## RESULTS

To test whether *Cbp53E* was involved in vision, we performed ERG recordings on *Cbp53E* null mutant flies. Two independent transgenic fly lines were tested: *Cbp53E*^*mi22*^ and *Cbp53E*^*mi41*^ (Hagel et al., 2015)). *Cbp53E* ^*mi22*^ and *Cbp53E*^*mi41*^ null mutants exhibited slower repolarization compared to *w*^*1118*^ controls following termination of the light stimulus (Figure 1). To attempt to quantify this difference, we measured the time to reach 75% repolarization after the light stimulus ended (see methods for details). Control *w*^*1118*^ flies took ∼0.5 seconds to repolarize, whereas *Cbp53E* null mutants took ∼1.5 seconds on average, a 3-fold increase in time needed to reach the 75% repolarization value (Figure 1). We compared other aspects of the ERG traces to see whether any of these were affected. *Cbp53E* ^*mi22*^ and *Cbp53E*^*mi41*^ null mutants displayed normal sustained negative potentials (SNP), ON-transients, and OFF-transients (Figure 2). We also compared ERG responses to tetanic stimulation, a protocol using short, rapid bursts of light to characterize temporal resolution (Krans et al., 2006). There were no obvious differences in the temporal dynamics of ERG responses to tetanic stimulation (Figure 2). These data indicate that loss of *Cbp53E* leads to delayed repolarization of photoreceptor cells following light stimulation, whereas activation of the phototransduction cascade and synaptic function appear to be normal.

Because calcium perturbation has been shown to lead to retinal degeneration in some cases, we wanted to test whether *Cbp53E* null exhibited age-related changes in ERG responses reflecting retinal degeneration (Wang & Montell, 2007a). We compared flies under 16-hour/8-hour light/dark cycles for up to 14 days. 14-day-old *Cbp53E*^*mi22*^ null mutants did not appear to have any age-dependent, or light-dependent changes to their ERG responses compared to 1 day old flies (Figure 3). *Cbp53E*^*mi22*^ null mutants also had normal rhodopsin levels upon eclosion (Figure 3). Another method that can be used to identify early stages of retinal degeneration is the ability to produce a prolonged depolarizing afterpotential (PDA), following stimulation with 480 nm wavelength blue light (Minke et al., 1975; Scavarda et al., 1983). *Cbp53E*^*mi22*^ flies exhibit normal PDA, even after 5 days of constant blue light exposure (Figure 3).

To see if we could rescue the mutant phenotype, we used the Gal4/UAS system to misexpress Drosophila Cbp53E in a *Cbp53E*^*mi22*^ null mutant background (Brand & Perrimon, 1993; Hagel et al., 2015). Because perturbation of either photoreceptor cells or supporting pigment cells can both lead to aberrant ERG responses, we performed cell-specific rescues using different GAL4 drivers (Wang & Montell, 2007b; Wang et al., 2010, 2012). We utilized either *rdhb-GAL4* to drive expression in pigment cells or *ninaE-GAL4* to drive expression in photoreceptor cells (O’Tousa et al., 1985; Wang et al., 2012). Expression of *UAS-Cbp53E* in photoreceptor cells, but not pigment cells, was able to rescue the *Cbp53E*^*mi22*^ null mutant phenotype, restoring the repolarization speed to rates that were similar to those of control *w*^*1118*^ flies (Figure 4). However, due to the variability inherent in the *Cbp53E*^*mi22*^ repolarization times, none of the rescue results were statistically significant compared to the *ninaE-GAL4* driver alone in the *Cbp53E*^*mi22*^ mutant background (Figure 4).

The closest human homologs to Drosophila Cbp53E are calbindin 2 (synonyms: CALB2, calretinin, CR, CB-D29k, CAL2, D29) and calbindin 1 (synonyms: CALB1, CAB1, D-28k) (Reifegerste et al., 1993a). We generated transgenic flies carrying *UAS-Calb2* and *UAS-Calb1* transgenes in a *Cbp53E*^*mi22*^ background to test whether the human homologs would rescue the drosophila ERG delayed repolarization phenotype. We tested two independent lines for the Cb2 rescue, and our only line for the Cb1 rescue, and found that all human calbindin homologs tested were able to rescue the mutant phenotype (Figure 4).

CALB2, CALB1 and other protein family members with six EF-hand domains have also been shown to function as calcium sensors that can participate in calcium-dependent signaling (Schwaller, 2020). We wondered whether Cbp53E was functioning as a calcium-sensor or simply as a calcium-buffer in our rescue experiments. To determine whether other Ca^2+^-buffering proteins, such as the smaller, two EF-hand containing protein, S100G (synonyms: calbindin-D9k, CB-D9k), would also rescue the *Cbp53E*^*mi22*^ null mutant phenotype, we generated *UAS-S100G* transgenic flies (see methods for details). Expression of S100G in a *Cbp53E*^*mi22*^ null mutant background yielded similar results and rescued the repolarization phenotype (Figure 4). This result suggests that the ability of Cb1, Cb2, and S100G to rescue the mutant phenotype when misexpressed in photoreceptor cells is most likely due to their calcium-buffering ability, rather than calcium-sensing and calcium-dependent signaling. While misexpression of all three of the human homologs tested was able to rescue the repolarization phenotype to levels indistinguishable from *w*^*1118*^ flies, none of these were found to be significantly different from the *ninaE-GAL4* driver alone, due to the high degree of variability in the repolarization times in the *Cbp53E* mutant background (Figure 4).

## DISCUSSION

We have demonstrated here that loss of Cbp53E leads to a delayed repolarization of photoreceptor cells in Drosophila. This phenotype is consistent in two independent mutant fly lines and can be rescued by expressing Cbp53E in photoreceptor cells using the GAL4/UAS system. This rescue can also be achieved by expressing the closest human homologs, respectively, CALB2 & CALB1. Our results suggest that loss of Cbp53E leads to a calcium imbalance within Drosophila photoreceptor cells, which contributes to the dynamics of neural responses to light stimuli *in vivo*. These results do not imply that Cbp53E is normally expressed in photoreceptor cells, as this has not been observed in other attempts to characterize its expression pattern (Reifegerste et al., 1993b). While we cannot rule out the possibility that Cbp53E is expressed in photoreceptor cells but was not detected previously, there are other potential explanations for the phenomena observed here, such as imbalances in extracellular calcium more generally as a result of Cbp53E loss in the entire animal.

The many roles of Ca^2+^ in Drosophila photoreceptor cells include activation, deactivation, and adaptation (Voolstra & Huber, 2020). Ca^2+^ perturbations have been shown to underlie many aspects of phototransduction in Drosophila, and exert many of these effects via the Ca^2+^-sensing protein, Calmodulin (CaM) (Voolstra & Huber, 2020). Mutations affecting either CaM or its target effectors often results in a prolonged deactivation phenotype (Scott et al., 1997; Smith et al., 1991). Furthermore, the slow termination phenotype we observe here is similar not only to that of Drosophila CaM mutants, but are also reminiscent of ERG phenotypes affecting *arr2* (major arrestin), *dcamta* (CaM-binding transcription factor), *inaC* (protein kinase C), *ninaC* (myosin III), and *rdgC* (Rhodopsin phosphatase), *norpA* (PLCβ) and its regulator, *stops* (SOCS box protein), all of which are closely involved in either regulating Ca^2+^ levels or responding to changes in Ca^2+^ levels in some way ((Wang et al., 2008) and reviewed in (Wang & Montell, 2007b) and (Voolstra & Huber, 2020)). We also note that in all of the photoreceptor-specific rescues, the shape of the SNP component of the ERG was similar to that seen in flies lacking *calx*, the calcium-sodium exchanger (Wang et al., 2005). There are still other calcium-binding proteins, such as the Ca^2+^ buffer protein expressed in photoreceptor cells, *calphotin*, was shown to lead to retinal degeneration in Drosophila when mutated (Weiss et al., 2012). Disruption of Cbp53E may affect other aspects of photoreceptor cell signaling. The human homologs, CALB2, CALB1, and secretagogin (SGN) likely function as Ca^2+^-sensors involved in signaling and modulation of cellular pathways (Schwaller, 2020). Our results suggest that the repolarization defects resulting from the loss of Cbp53E could be rescued by misexpression of calcium-buffering proteins, whether containing 6 EF-hand domains (CALB1, CALB2) or 2 EF-hand domains (S100G).

## ACKNOWLEDGEMENTS

Research reported in this publication was supported by an Institutional Development Award (IDeA) from the National Institute of General Medical Sciences (NIGMS) of the National Institutes of Health (NIH) under grant number P20GM103442. We thank Dr. Charles Tessier for the kind gift of Cbp53E mutant fly stocks. We also wish to thank Drs. Daniel Barr, Joseph Biggane, John H. Boyle, Wendy Larson, and James Peliska for critical feedback and helpful suggestions related to this manuscript.

